# Transcriptome profiles of *T.b. rhodesiense* in Malawi reveal focus specific gene expression Profiles associated with pathology

**DOI:** 10.1101/2023.07.11.548495

**Authors:** Peter Nambala, Harry Noyes, Joyce Namulondo, Oscar Nyangiri, Enock Matovu, Vincent Pius Alibu, Barbara Nerima, Annette MacLeod, Janelisa Musaya, Julius Mulindwa, the TrypanoGEN+ Research Group as Members of the H3Africa Consortium

## Abstract

**Background:** Sleeping sickness caused by *T.b. rhodesiense* is a fatal disease and endemic in Southern and Eastern Africa. There is an urgent need to develop novel diagnostic and control tools in order to achieve elimination of rhodesiense sleeping sickness which might be achieved through a better understanding of trypanosome gene expression and genetics using endemic isolates. Here, we describe transcriptome profiles and population structure of endemic *T. b. rhodesiense* isolates in human blood in Malawi.

**Methodology:** Blood samples of r-HAT cases from Nkhotakota and Rumphi foci were collected in PaxGene tubes for RNA extraction before initiation of r-HAT treatment. 100 million reads were obtained per sample, reads were initially mapped to the human genome reference GRCh38 using HiSat2 and then the unmapped reads were mapped against *Trypanosoma brucei* reference transcriptome (TriTrypDB54_TbruceiTREU927) using HiSat2. Differential gene expression analysis was done using the DeSeq2 package in R. SNPs calling from reads that were mapped to the *T. brucei* genome was done using GATK in order to identify *T.b. rhodesiense* population structure.

**Results:** 24 samples were collected from r-HAT cases of which 8 were from Rumphi and 16 from Nkhotakota foci. The isolates from Nkhotakota were enriched with transcripts for cell cycle arrest and stumpy form markers, whereas isolates in Rumphi focus were enriched with transcripts for folate biosynthesis and antigenic variation pathways. These parasite focus-specific transcriptome profiles are consistent with the more virulent disease observed in Rumphi and a more silent disease in Nkhotakota associated with the non-dividing stumpy form. Interestingly, the Malawi *T.b. rhodesiense* isolates expressed genes enriched for reduced cell proliferation compared to the Uganda *T.b. rhodesiense* isolates. PCA analysis using SNPs called from the RNAseq data showed that *T. b. rhodesiense* parasites from Nkhotakota are genetically distinct from those collected in Rumphi.

**Conclusion:** Our results have added new insights on how clinical phenotypes of r-HAT in Malawi might be associated with differences in gene expression profiles and population structure of *T*. *b. rhodesiense* from its two major endemic foci of Rumphi and Nkhotakota.

**Author Summary:** A better understanding of *T. b. rhodesiense* gene expression profiles and population structure using endemic isolate may fast track the current search for novel diagnostic and control tools for rhodesiense sleeping sickness. Here, we analysed *T. b. rhodesiense* transcriptome profiles from endemic isolated from peripheral blood in Nkhotakota and Rumphi foci in Malawi. In Nkhotakota focus, *T. b. rhodesiense* transcripts were enriched for cell cycle arrest and stumpy marker whereas in Rumphi focus, the isolates were enriched for antigenic variation and folate biosynthesis biological pathways. Furthermore, we also found that *T. b. rhodesiense* population structure in Nkhotakota focus is different from Rumphi focus. The differences in trypanosome gene expression profiles and population structure are consistent with a less severe and acute sleeping sickness clinical profiles in Nkhotakota and Rumphi foci respectively.

## Introduction

Human African trypanosomiasis (HAT) causes social and economic burdens on people living in remote areas where HAT is endemic. *Trypanosoma brucei gambiense* is the causative agent of gambiense HAT (g-HAT) in Western, Central and part of Eastern Africa, whereas, *Trypanosoma brucei rhodesiense* (Tbr) causes rhodesiense HAT (r-HAT) in Southern and Eastern Africa. Although, intensified HAT surveillance championed by the World Health Organisation has led to drastic decreases in HAT incidences over the past 20 years, r-HAT is still endemic in Malawi in areas adjacent to wildlife reserves such as Nkhotakota and Rumphi districts (1). For example, there was a sudden surge in r-HAT cases from 2019 to 2021 in Malawi’s r-HAT foci and the cause is yet to be established (2). To prevent future episodes of HAT outbreaks, there is need to develop novel epidemiological interventions and surveillance tools which might be achieved through a comprehensive understanding of the parasite genetics and gene expression profiles underlying HAT transmission cycle.

Genetic recombination between *T. b. rhodesiense* and *T. brucei brucei* during circulation in animals and tsetse vectors is believed to result in creation of new strains that may impact the epidemiological landscape of HAT diseases as it may create a genetic pool of human infective trypanosomes (3). Genetic characterisation of Tbr isolates from Malawi and Uganda revealed that the genetic structure of trypanosomes between the two countries is different with *T. b. rhodesiense* isolates in Uganda being more clonal compared to Malawi isolates that had greater genetic diversity and evidence of frequent mating (4). Moreover, r-HAT in Uganda tends to be an acute disease whereas it ends to be a more chronic disease in Malawi’s Nkhotakota focus (5). At the same time, genetic diversity of *T. b. rhodesiense* isolates have been observed between Uganda’s three r-HAT foci with expansion of new foci from Kenya (6). It remains unclear to what extent differences in clinical presentation are associated with parasite or host genetic diversity. We previously described human transcriptome profiles in r-HAT disease in Nkhotakota and Rumphi foci in Malawi where r-HAT clinical presentation is different between the two foci (7). We also found that there were differences in expression profiles in individuals with stage 1 and stage 2 disease but no differences in infected individuals between Nkhotakota and Rumphi foci (7).

Therefore, the current study describes transcriptome profiles and population structure of T.b. rhodesiense parasites isolated from r-HAT patients in the endemic foci of Nkhotakota and Rumphi in Malawi. Additionally, we have also compared the transcriptome profiles of Tbr isolates between Malawi and Uganda. Our data shows that the differences in pathology between the two foci is associated with differences in parasite population structure. This may contribute to the search for novel trypanosome diagnostic markers and control strategies.

## Methods

### Ethics statement and Sample Collection

The r-HAT surveillance and study participants recruitment for blood sample collection to be used for dual RNA sequencing has been previously described (2). Ethical approval of the study was obtained from the Malawi National Health Sciences Research Committee (Protocol Number: 19/03/2248). Consent and assent were obtained from each study participant before sample collection. Briefly, sample collection was done during active and passive r-HAT surveillances conducted for 18 months from July 2019 to December 2020. HAT cases were confirmed to be infected with trypanosome parasites by microscopic examination of thick blood films during the surveillance period. 2ml whole blood samples were collected into Paxgene^®^ tubes from r-HAT cases and stored at -20°C until processing. All samples were collected before initiation of r-HAT treatment and all patients were thereafter treated following the national r-HAT treatment guidelines. Thereafter, a PCR targeting the serum resistance associated (SRA) gene of *T. b. rhodesiense* (28), was used to confirm rhodesiense HAT disease in recruited study participants.

### RNA sequencing and analysis

Dual RNA sequencing was done on the same samples that we used for human transcriptome analysis we previously described. Since trypanosomes are blood parasites it was possible to obtain trypanosome transcriptomes from the same RNA-seq data. Briefly, RNA was extracted from blood of *T. b. rhodesiense* infected individuals using TRIzol method (29). Samples with total RNA >1µg were selected for RNA library preparation using the QIASeq FastSelect rRNA, globin mRNA was depleted and libraries were prepared for sequencing Illumina NovaSeq with the <u>NEBNext Ultra II Directional RNA Library Prep Kit</u> a target depth of 100 million reads. FASTQ reads were aligned to the GRCh38 release 84 human genome sequence obtained from Ensembl (Howe et al., 2021) using HiSat2 (Kim et al 2019). Unmapped reads were then mapped against reference transcriptome TriTrypDB54_TbruceiTREU927 using HiSat2. After mapping to the *T. brucei* transcript sequence only reads that were mapped in proper pairs were retained in the output bam-file for differential gene expression in R Studio V4.2 using DESeq2 package (30), transcripts with less than 10 reads across all samples were filtered out. Enriched biological pathways were determined by uploading significant (padj<0.05) differentially expressed genes in TriTryDB release 61 (13).

### *T.b. rhodesiense* SNP Calling and analysis

The GATK workflow was used for SNP calling from reads that were mapped to the *T.brucei* transcriptome. VSG and ESAG genes were removed prior to population structure analysis since the numerous copies of these genes make mapping difficult, leading to SNP calls due to miss-alignment rather than mutation (31). The resulting SNP data set was used in PLINK line for: i) multidimensional scaling (MDS) analysis based on raw Hamming genetic distance to generate principal coordinates as well as population cluster distance matrix and ii) for Fixation index estimation of allele variance based on Wright’s F-Statistics (*F_ST_*) (21, 32). Graphical output of PCA and hierarchical clustering was prepared in R Studio version 4.3.1(33).

## Results

### No Differences in *T.b. rhodesiense* Transcriptomes from Stage 1 and 2 HAT isolates

We had previously identified differences in human transcriptomes between individuals with stage 1 and stage 2 HAT disease in Malawi (7). We used RNA-seq data from the same samples of r-HAT cases to align to the *T. brucei* transcriptome for trypanosome transcriptome analysis. A total of 16 and 8 cases were collected from Nkhotakota and Rumphi respectively. We compared transcriptomes of *T.b. rhodesiense* isolated in individuals with stage 1 and 2 HAT, no *T.b. rhodesiense* genes were significant differentially expressed (DE) between these two stages. There was also no difference in gene expression of *T.b. rhodesiense* parasites isolated from male and female HAT cases.

### Enrichment of Cell Cycle Arrest and Stumpy form Marker Transcripts in Nkhotakota *T.b. rhodesiense* isolates

Clinical presentation of HAT disease in Malawi is focus dependent (2). To determine the contribution of variations in trypanosome gene expression to focus specific clinical phenotypes, we performed a principal component analysis (PCA) and differential transcriptome analysis of *T.b. rhodesiense* genes of parasites from Nkhotakota and Rumphi focus. Four samples were identified as outliers in the PCA and were excluded from the analysis (**Fig S1A**). Principal components 2 and 3 separated *T.b. rhodesiense* transcriptomes from the two HAT foci into two distinct clusters **(Fig 1A)**, and a total of 91/10628 (0.86 %) genes were significant (padj < 0.05) differentially expressed between the two foci. Of the 91 genes, 21/91 (23.08 %) genes were upregulated log2 fold change (log2FC) >1 in isolates from Nkhotakota (**Fig 1B and Table S1**), of which 43 % (9/21) of the genes encode for ESAG1, ESAG9 and ESAG11. ESAG1 is a *T. brucei* heat shock protein that induces differentiation of procyclic trypanosomes to blood stream forms *in vitro* when there is a temperature shift from 27°C to 37°C (8). ESAG 9 is a stumpy form marker that is enriched in *T. brucei* as adaptation for either tsetse vector transmission or for sustainability of infection chronicity in mammalian hosts (9, 10). ESAG11 encodes GPI transmembrane protein which plays a role in lipid raft and glycosylation of *T. brucei* VSGs (11, 12). Two DEGs encoded VSG **(Fig S1B)**.

**Figure 1.**
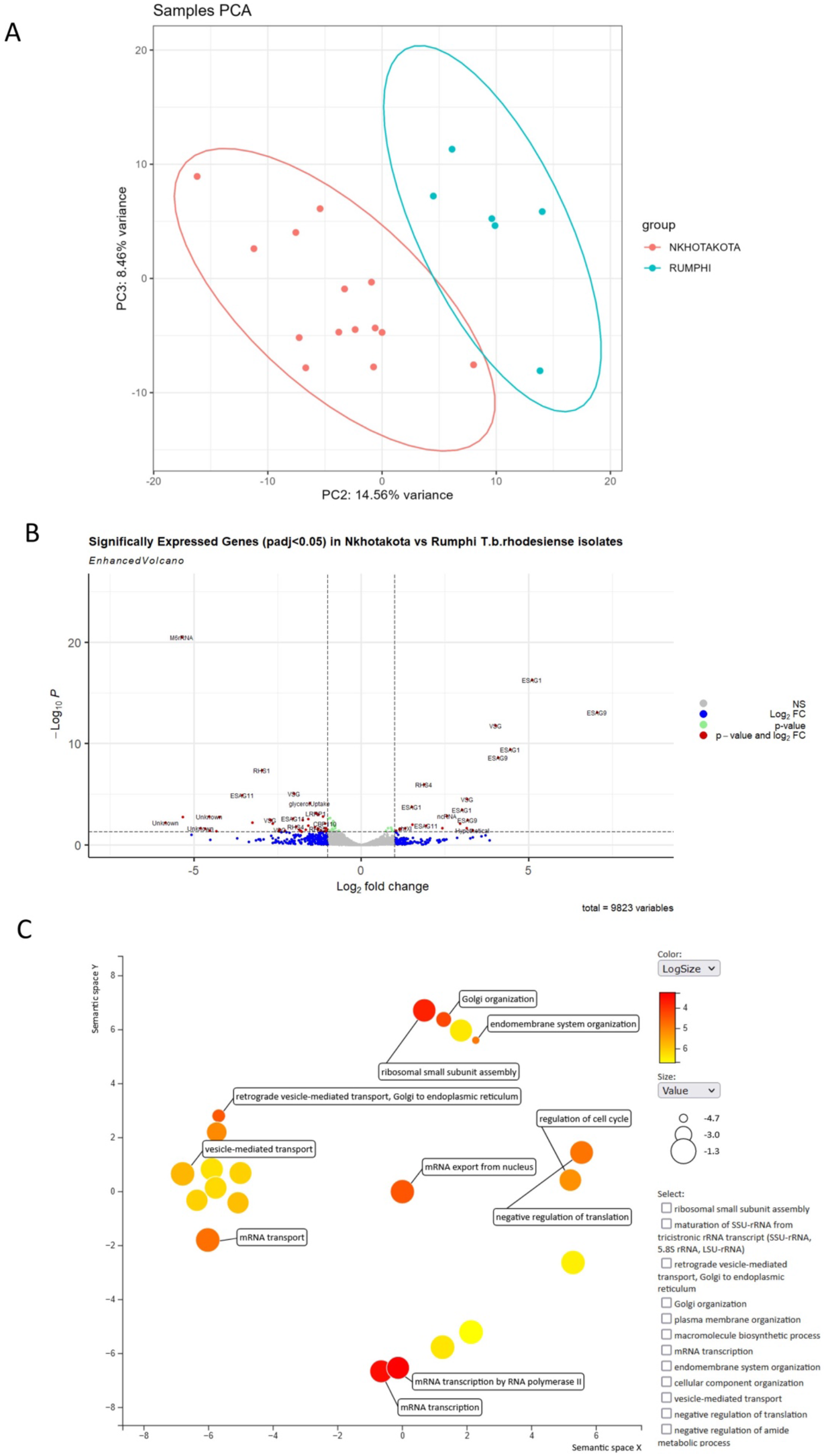
Differentially expressed *T.b. rhodesiense* genes. **A)** stratification of *T.b. rhodesiense* transcriptomes in isolates between Nkhotakota and Rumphi focus on a plot of PC2 and PC3. **B)** Genes that were upregulated with log2FC >1 in isolates from Nkhotakota. **C)** Biological pathways of *T.b. rhodesiense* upregulated genes enriched in during human infection in Nkhotakota focus. The axex in the plot have no intrinsic meaning but semantically similar GO terms remain together in the plot (16).

To determine biological pathways exploited by *T.b. rhodesiense* during human infection in Nkhotakota focus, upregulated genes (21/91) were loaded into TriTrypDB release 60 (13). For this, we observed genes that were enriched (p<9.63E-3) for endomembrane system organization, retrograde vesicle-mediated transport, Golgi organization, Golgi vesicle transport and plasma membrane organization **(Fig 1C**, **Fig S2 and Table S2)**. We also observed enrichment of genes for mitotic cell cycle arrest, negative regulation of cellular amide metabolic process, negative regulation of cellular protein metabolic process and negative regulation of protein metabolic process.

**Figure 2.**
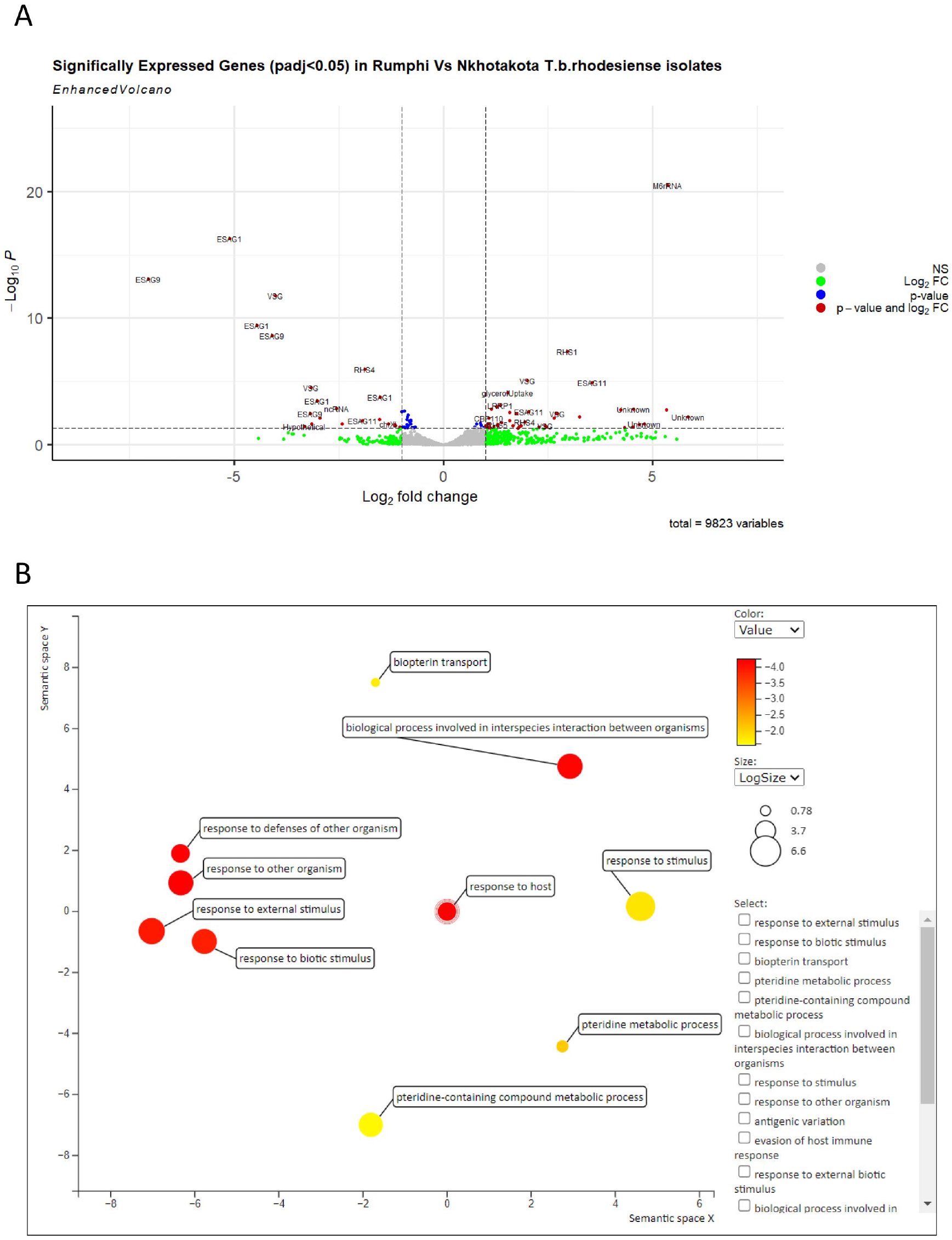
Differential gene expression of Tbr isolates from Rumphi focus. **A)** Genes that were upregulated with log2FC >1. **B)** Biological pathways of *T.b. rhodesiense* upregulated genes enriched in during human infection. Folate biosynthesis and response to host were among the enriched pathways.

Collectively, these results indicate that the *Tbr* isolates from Nkhotakota were predominantly in the stumpy form which could explain the chronic form of HAT in this focus when compared to Rumphi.

### Enrichment of folate biosynthesis and antigenic variation transcripts in Rumphi *T.b. rhodesiense* Isolates

From the 91/10628 differentially expressed genes (DEGs) in *T.b. rhodesiense* isolates, 43/91 (47.25%) genes were upregulated (padj> 1.0) in isolates from Rumphi focus **(Fig 2A and Table S1)**. Of the 43 genes, 14 (33%) coded for VSGs (log2FC >1.7-5.8), and4 (9%) for ESAGs. M6 rRNA was the most significant (padj<3.35E -25) differentially expressed gene and highly upregulated (log2FC >5) suggesting high protein synthesis in Rumphi *Tbr* isolates compared to isolates in Nkhotakota focus. Kinesin K39 was also upregulated in Rumphi parasites and is an ATP-dependent cytoskeleton motor protein which in eukaryotic cells plays a crucial role in cell cycle and migration (14). In *Leishmania donovani*, a trypanosomatid that causes leishmaniasis, K39 kinesis accumulates and moves along the cortical cytoskeleton in a cell cycle-dependent preference for the posterior pole of the cell (15).

Next, we uploaded the 43 DEGs upregulated in Rumphi focus into TriTrypDB release 60 (13), to determine biological processes that were enriched and visualised the enriched biological processes in REVIGO (16). For this we identified high enrichment (48.19 to 144.58 fold enrichment) of Pteridine metabolic processes and transport **(Table S3)**. Pteridine together with folic acid are essential folates used for metabolic biosynthesis of DNA, RNA and amino acids

(17). Trypanomastids exploits pteridine and folic acid metabolites in mammalian hosts and insect vectors for folate biosynthesis of purine and pyrimidine nucleotides. We also observed enrichment of *T. b. rhodesiense* biological processes involved in response and evasion of host immune response as well as in pathogen-host interaction **(Fig 2B)**. These results suggest that the Tbr isolates from Rumphi were enriched for bloodstream forms that are highly replicative and exploiting the human folate metabolites for nucleotide synthesis. This could perhaps explain the observed acute nature of rHAT disease in Rhumpi as compared to Nkhotakota.

### Malawi *T. b. rhodesiense* parasites are enriched with cell cycle arrest transcripts compared to Uganda *T. b. rhodesiense* parasites

Since *T. b. rhodesiense* gene expression is different between Malawi’s r-HAT foci and clinical presentation of r-HAT varies between countries (5, 18), we next sought to compare gene expression of Malawi parasites with published data from Ugandan parasites (19, 20). We first identified gene IDs that were mapped to both Malawi and Uganda *Tbr* isolates and filtered out gene IDs found in isolates of one country only. In total, 7003 *Tbr* Gene IDs had counts that were common to both Malawi and Uganda isolates were then loaded into DEseq2 for differential gene expression and principal component analysis. There was a distinct clustering in PCA1 and PCA3 between Malawi and Uganda human *Tbr* isolates **(Fig S3).** Parasites from Uganda also clustered together and were distinct from the Malawi parasites suggesting clonality of isolates, and isolates from rodents clustered differently from human isolates of both countries. Since rodents samples clustered differently from human samples, we removed them from further expression analysis.

**Figure 3.**
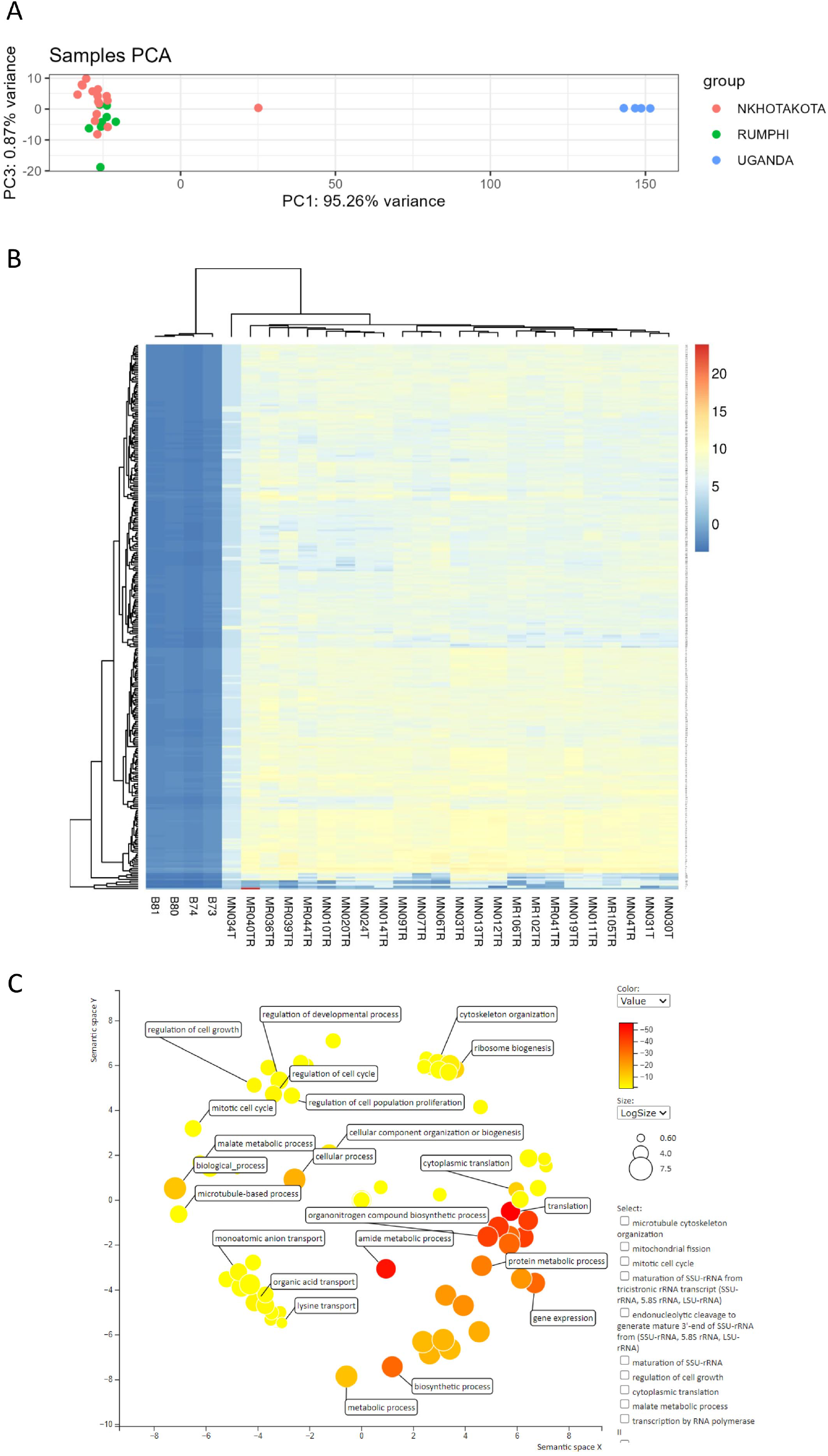
Comparison of gene expression profiles between Malawi and Uganda Tbr isolates. **A)** PCA analysis of Tbr isolated in individuals with stage 1 and stage 2 r-HAT in Malawi and Uganda. Sample MN034T was intermediate between Malawi and Uganda Tbr isolates **B)** A eucledian heatmap generated in PCAExploler comparing the gene explession level of each Tbr isolate from Malawi and Uganda. Sample MN034T that was intermediate in PCA also had an intermediate gene expression levels compared to other Malawi isolates **C)** Enriched biological pathways of Malawi Tbr isolates compare to Uganda isoltes loaded in TriTrypDB release 60 and visualised in REVIGO (13, 16). Most of the transcripts were enriched for reduced cell proliferation in Malawi isolates compared to Uganda isolates.

A comparison of human Tbr isolates from Malawi and Uganda showed that 3132/7003 (44.72%) gene were significantly (padj< 0.05) differentially expressed of which 1565/3132 (49.97%) gene were upregulated in Malawi (log2FC >1) and 753/3132 (24.04%) genes were downregulated (**Fig 3A and 3B)**. We further identified 127/1565 (8.12%) genes that were significant differentially expressed with padj< 8.33E -35 and highly upregulated (log2FC > 13.5 to 17.0) in Malawi Tbr isolates. Among the most upregulated genes were *T.brucei* protein Associated with Differentiation (TbPAD2, log2FC 17.0) and TbPAD2 (log2FC 16.0) which are stumpy markers for blood stream trypanosomes. To identify gene ontology biological pathways enriched by the 127 genes, we uploaded the gene list in TriTrypDB release 60 and visualised in REVIGO (13). This identified regulation of cell growth, regulation of growth, mitotic cell cycle arrest, regulation of cell population proliferation and regulation of developmental process among the enriched biological pathways for Malawi *T. b. rhodesiense* isolates compared to Uganda parasites **(Fig 3C)**. This suggests that Malawi *T. b. rhodesiense* are enriched with stumpy parasites which are necessary for efficient transmission in tsetse vector compared to Uganda *T. b. rhodesiense* and could explain the more chronic nature of the Malawi strain as compared to the Uganda strain.

### Population Structure and Genetic diversity of *T. b. rhodesiense* isolates Varies Between Rumphi and Nkhotakota Foci

Having identified that *T. b. rhodesiense* gene expression profiles in Malawi are focus specific, we next sought to understand whether there are differences in allele frequencies between Nkhotakota and Rumphi isolates. SNP calling from reads that were mapped to the *T.brucei* genome was done using GATK workflow and loaded in PLINK command line for multidimensional scaling (MDS) analysis based on raw Hamming genetic distance (21). The results showed a clear population stratification and genetic distance of isolates from Nkhotakota and Rumphi focus on principal components (PC) 1 and PC 2 **(Fig 4A)**. The distance matrix was then used to construct a phylogenetic tree with unrooted neighbour-joining without assuming evolutionary hierarchy. The phylogenetic tree showed a clear genetic distance between *T. b. rhodesiense* populations in Nkhotakota and Rumphi focus **(Fig 4B)**. We next used fixation index (Fst) to measure the between groups genetic variance in *T. b. rhodesiense* populations from Nkhotakota and Rumphi. Using many SNP markers called from RNA-seq data, it is possible to get an estimate of genetic differentiation without needing to use a large sample size (22). An Fst value of 1 suggests complete differentiation in allele frequency between subpopulations while a value of 0 suggests no differentiation and a mean Fst of greater than 0.15 is considered significant in differentiating populations (21). There was a mean Fst of 0.31 between the two populations and 352 SNPs had an Fst of 1.0 of which most were on chromosome 1 **(Fig 4C).** The results suggest diversity in *Tbr* population structure between Nkhotakota and Rumphi focus isolates which might be due to fixation of alleles in one population or the other.

**Figure 4.**
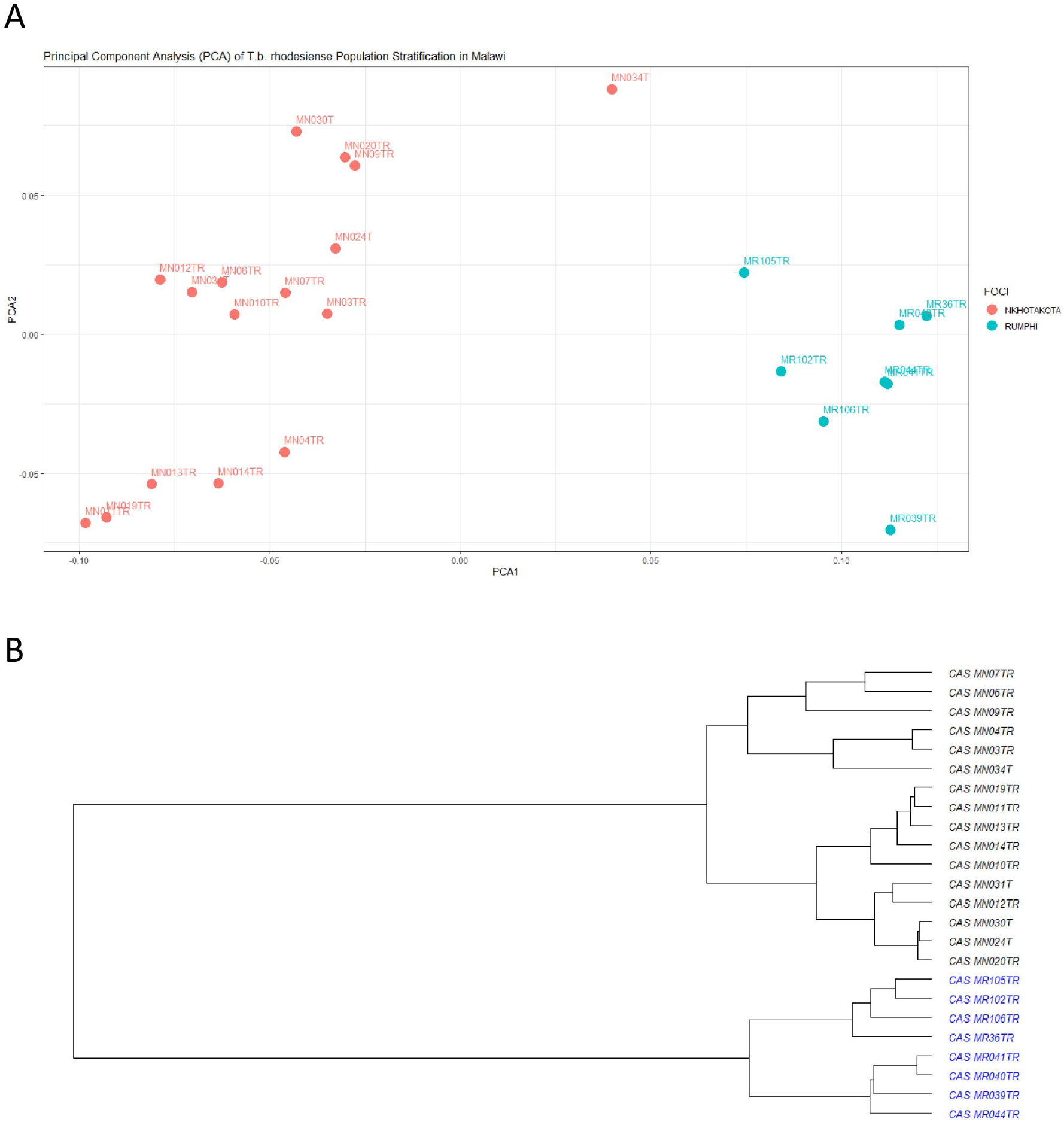
Polulation structure and genetic diversity of Tbr between Nkhotakota and Rumphi foci. **A)** population stratification and genetic distance of isolates from Nkhotakota and Rumphi foci on Principal Component Analysis (PCA) 1 and PCoA 2. **B** Unscaled hierarchical clustering dendogram showing the relatedness of Tbr isolates from Nkhotakota and Rumphi foci that was generated using population cluster distance matrix. Blue and black color represents isolates from Rumphi and Nkhotakota respectively

## Discussion

In the current study we have compared gene expression profiles and population structure of *T. b. rhodesiense* isolates between Nkhotakota and Rumphi foci. Additionally, we have compared gene expression profiles between Malawi and Uganda, representing Southern Africa and Eastern Africa respectively. Overexpression of VSGs in Rumphi isolates suggest high proliferation of bloodstream slender form trypanosomes compared to isolates in Nkhotakota. Slender Trypanosomes use a repertoire of VSGs to evade host adaptive immune system whereas, stumpy trypanosomes do not express VSGs. This is consistent with differential expression of kinesin K39 which maintains cell cytoskeleton integrity during the cell cycle. Moreover, high human antibody titre against *Leishmania chagas* and *Leishmania donovani* kinesin K39 antigen has been detected in patients with Chagas disease making kinesin K39 a potential biomarker for serological diagnosis (23). Identification of kinesin K39 in *T. b. rhodesiense* isolates highlights this as a potential biomarker for a much need serodiagnosis of *T. b. rhodesiense* infections and should be validated in future studies.

We have also identified high enrichment of transcripts for pteridine in *T. b. rhodesiense* isolates form Rumphi focus. Structural differences between the fusion protein DHFR-ThyS in trypanosomatids and the individual polypeptides in humans make this protein (folate) an attractive target for rational drug design which should be exploited in future research. Additionally, exploitation of host folate metabolites by *T. b. rhodesiense* isolate may have implications of clinical pathology of r-HAT as it may induce host anaemia when the parasite uses haemoglobin as a folate source. Consistent with this finding, studies on clinical presentation of HAT in Malawi had identified anaemia as one of the clinical pathology associated with HAT patients (24).

The contrasting enrichment of genes required for adaption of *T. b. rhodesiense* transmission to tsetse fly vector and sustainability of blood stream slender trypanosomes in Nkhotakota and Rumphi foci may have implications on r-HAT clinical phenotype, control and elimination. Indeed, we previously established that most r-HAT cases in Nkhotakota present with a stage 1 disease whereas in Rumphi most r-HAT cases present with a severe stage 2 r-HAT disease (2). Virulence of trypanosome infection in mammalian host is determined by accumulation of a population of slender trypanosomes (25). We speculate that severe acute cases of r-HAT observed in Rumphi in comparison to Nkhotakota, might be due to the ability of isolates in Rumphi to maintain high population of slender trypanosomes whereas those in Nkhotakota foci have predominantly the non-dividing stumpy forms and hence lower parasitaeimia. We further propose that *T. b. rhodesiense* isolates in Nkhotakota focus might be highly transmissible as they overexpressed stumpy markers whereas isolates in Rumphi focus maybe less transmissible due to high maintenance of a slender trypanosome population during human infections. Nonetheless, future research should consider validating our current findings using appropriate experimental models.

Comparison of gene expression profiles between Malawi (Southern Africa) and Uganda (East Africa) showed distinct clustering of samples between the two countries which is consistent with microsatellite analysis results of isolates from Malawi and Uganda (4, 6). Additionally, previous population genetics analysis identified that Ugandan isolates have a clonal population compared to diversified Malawi (Nkhotakota focus) isolates which was consistent with our transcriptomics results. Enrichment of cell cycle arrest biological pathways in Malawi isolates demonstrates the need for control strategies to focus on breaking the contact cycle between humans and tsetse fly vectors in Malawi r-HAT foci. Stratification of samples from human and rodents also suggests that *T. brucei* exploits different genes when circulating in dissimilar mammalian hosts. Inference from animal models or cell culture results on disease in humans should be done with caution as it may not be a true representation of *T. b. rhodesiense* infection dynamic in humans. For example, all expressed VSG that were identified by Jayaraman et.al., through long read sequencing of *T. brucei* passages in mice (26), were not differentially expressed in our data and vice versa. This might explain the challenge that has been in identifying expressed antigens that may be used as novel biomarkers for diagnostics and vaccine development in humans.

We have also identified 2 and 14 unique VSGs in Nkhotakota and Rumphi isolates respectively, that were expressed by *T. b. rhodesiense* in all blood samples analysed. Blood samples used in the current study were randomly collected over a period of 18 months excluding the possibility that the expressed VSG were randomly expressed. Although most expressed VSGs are highly antigenic and constantly changing, some VSG also elicit little host antibody response thereby subverting natural immunity (27). The identified VSGs have a potential to be explored in future research to determine if they continue to be consistently expressed, then they could be used as biomarkers for a much-needed rapid diagnostic test and vaccine against *T. b. rhodesiense* infections. In animal trypanosomes, a unique VSG expressed throughout infection has been used to develop a vaccine candidate that offered protection against *T. vivax* infection in mice models which has a potential of a clinical trial (27).

In conclusion, our results from both gene expression profiles and population genetic analysis have added new insights on how clinical phenotypes of r-HAT might be influenced by differences in *T. b. rhodesiense* population structure and gene expression profiles. We have used RNA-seq data to call *T. b. rhodesiense* SNPs from endemic isolates which will contribute to future studies of *T. b. rhodesiense* population genetics. SNP analysis results showed a distinct stratification in the *T. b. rhodesiense* population structure between isolates from Nkhotakota and Rumphi foci suggesting that there is little mixing of parasites between these two foci and that there is potential to control the infection in each focus independently. This is consistent with the results we obtained from trypanosome gene expression profiles which showed distinct clustering of gene expression profiles of isolates form each focus. Additionally, we have showed that transcriptome profiles of *T. b. rhodesiense* isolates in Nkhotakota and Rumphi are different. Peripheral blood trypanosomes in Nkhotakota were enriched with transcripts for stumpy trypanosomes, whereas in Rumphi, the trypanosome transcripts were enriched for antigenic variation and folate biosynthesis. Lastly, we have also identified differences in transcriptome profiles between Malawi and Uganda *T. b. rhodesiense* isolates. Pulled transcriptomes of Malawi *T. b. rhodesiense* were enriched for cell cycle arrest compared to Uganda isolates. Future research should consider validating our findings by obtaining pathological markers in rodents infected with *T. b. rhodesiense* isolates from Nkhotakota and Rumphi foci.

## Author Contributions

**Peter Nambala:** Conceptualization, Methodology, Investigation, Formal analysis, Writing - original draft. **Harry Noyes:** Conceptualization, Methodology, Formal analysis, Writing - review & editing. **Vincent Pius Alibu:** Conceptualization, Writing - review & editing, Methodology. **Barbara Nerima:** Conceptualization, Writing - review & editing, Methodology. **Joyce Namulondo:** Formal analysis**. Oscar Nyangiri:** Formal analysis. **Enock Matovu:** Conceptualization, Supervision. **Annette MacLeod:** Conceptualization. **Janelisa Musaya:** Conceptualization, Writing - review & editing, Methodology, Supervision, Formal analysis. **Julius Mulindwa:** Conceptualization, Writing - review & editing, Methodology, Formal analysis, Supervision.

## Acknowledgement

We would like to acknowledge Nkhotakota and Rumphi district health offices for their assistance in sample collection.

## Supplementary data

**Figure S1.**
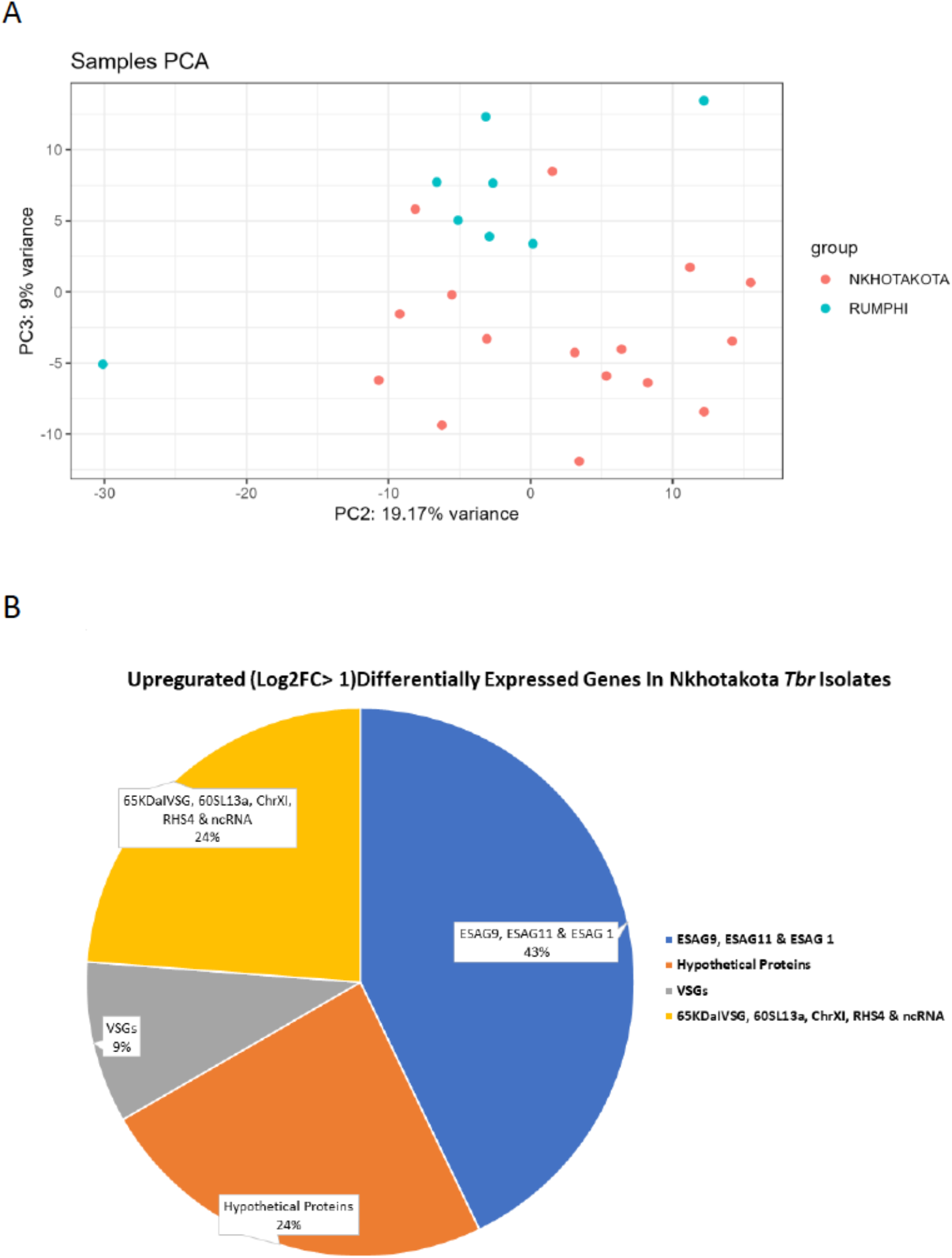
Differential gene expression analysis of Tbr isolates from Nkhotakota versus Rumphi foci. **A)** PCA analysis showing outlier samples from both Nkhotakota and Rumphi foci that were excluded from further analysis. **B)** Proportions of upregulated differentially expressed genes in in Nkhotakota Tbr isolates with ESAG transcripts being the most upregulated

**Figure S2.**
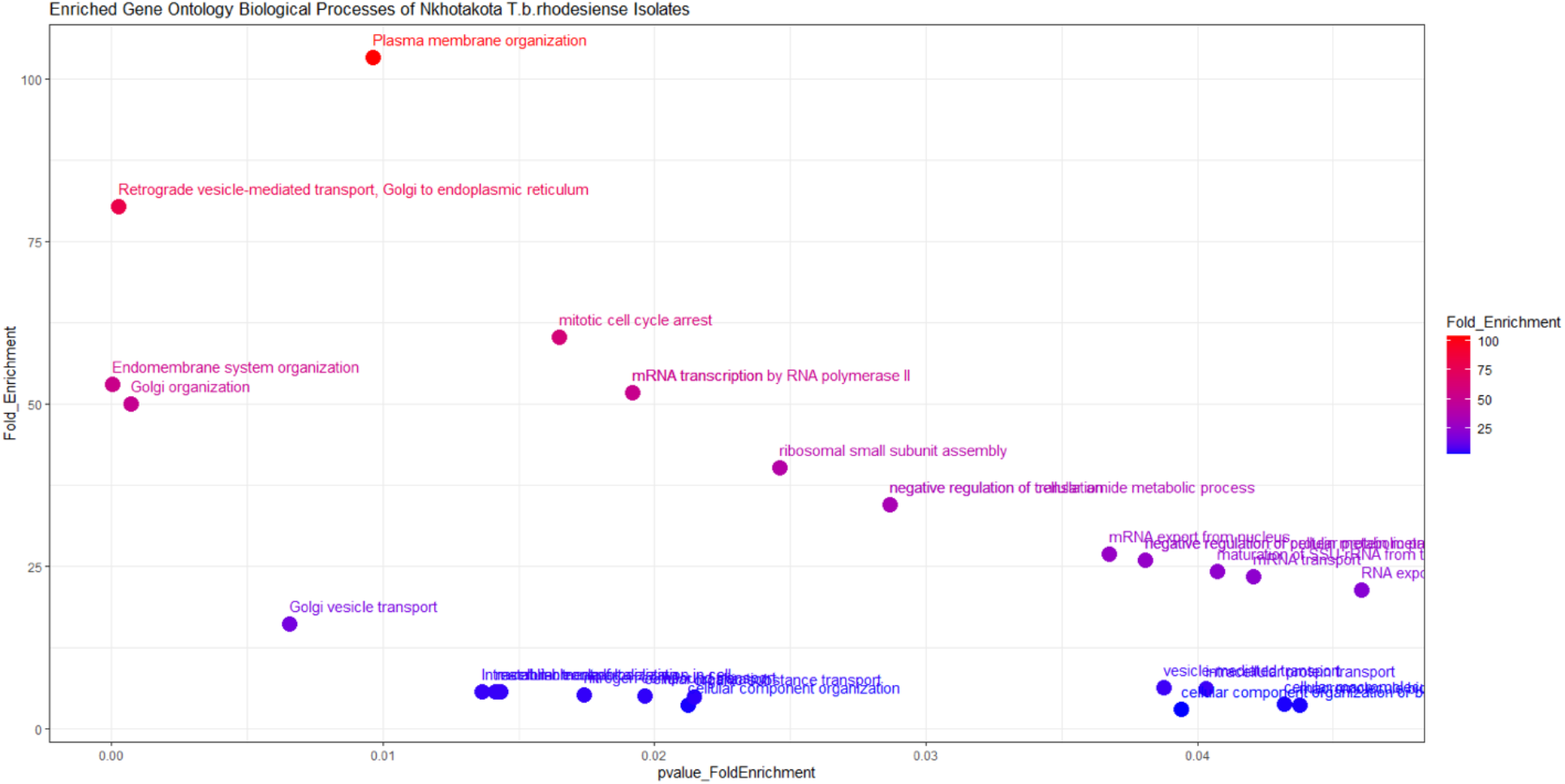
Fold enrichment of gene ontology biological processes of Nkhotakota Tbr isolates loaded in TriTrypDB (11).

**Figure S3.**
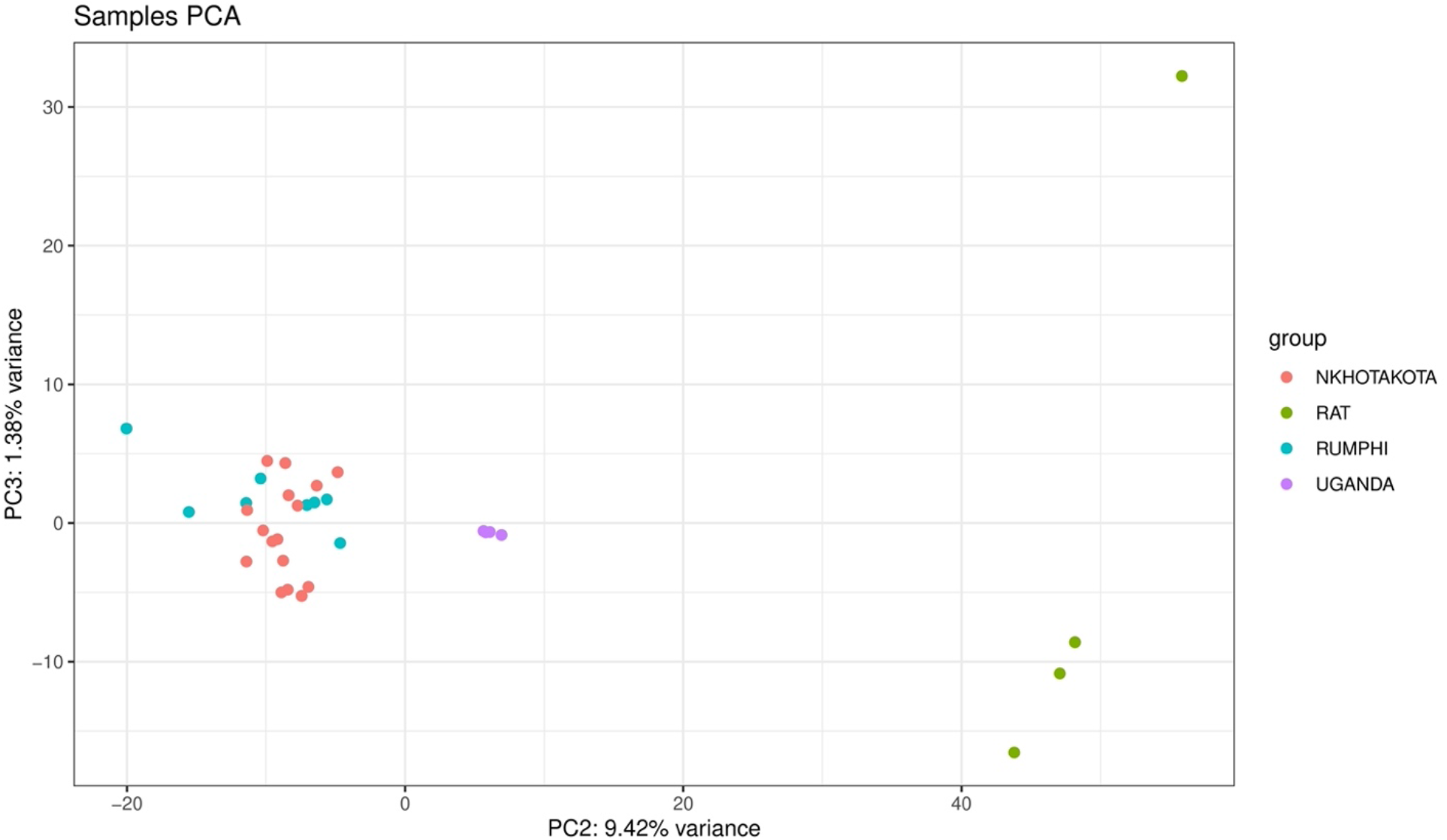
Principal component analysis of Tbr transcriptomes comparing human isolates from Nkhotakota (16 isolates), Rumphi (Eight isolates), Uganda (Four isolates) and Tbr isolates passaged in rodents.

**Table S1:** List of Genes that were significant (padj<0.05)differentially expressed in Tbr isolates from Nkhotakota focus versus Rumphi focus

| GeneID | GeneName | padj | Nkhota<br>log2FC | Rumphi<br>log2FC |
| --- | --- | --- | --- | --- |
| Tb927.1.5220:mRNA | expression site-associated gene 9 (ESAG9) protein | 8.571E-14 | 7.05 | -7.05 |
| Tb09.v4.0065:mRNA | expression site-associated gene 1 (ESAG 1) | 5.393E-17 | 5.11 | -5.11 |
| Tb927.11.17890:mRNA | expression site-associated gene 1 (ESAG1) protein | 3.933E-10 | 4.46 | -4.46 |
| Tb927.9.680:pseudogenic_transcript | expression site-associated gene 9 (ESAG9) | 2.434E-09 | 4.09 | -4.09 |
| Tb927.11.17870:mRNA | variant surface glycoprotein (VSG) | 1.619E-12 | 4.01 | -4.01 |
| Tb927.9.700:mRNA | hypothetical protein | 3.518E-02 | 3.32 | -3.32 |
| Tb927.11.18670.1 | expression site-associated gene 9 (ESAG9) | 3.542E-03 | 3.18 | -3.18 |
| Tb927.9.690:pseudogenic_transcript | variant surface glycoprotein (VSG) | 3.032E-05 | 3.17 | -3.17 |
| Tb11.1000:mRNA | expression site-associated gene 9 (ESAG9) protein | 2.298E-02 | 3.15 | -3.15 |
| Tb927.9.660:mRNA | expression site-associated gene 1 (ESAG1) protein | 3.397E-04 | 3.01 | -3.01 |
| Tb11.v5.0984.1 | hypothetical protein | 7.535E-03 | 2.95 | -2.95 |
| Tb9.NT.6:ncRNA | Noncoding RNA | 1.396E-03 | 2.56 | -2.56 |
| Tb927.5.150:mRNA | hypothetical protein | 2.298E-02 | 2.43 | -2.43 |
| Tb927.1.4900:mRNA | expression site-associated gene 11 (ESAG11) protein | 1.320E-02 | 1.93 | -1.93 |
| Tb927.2.510:mRNA | retrotransposon hot spot protein 4 (RHS4) | 1.103E-06 | 1.88 | -1.88 |
| Tb927.7.370:mRNA | hypothetical protein | 9.459E-03 | 1.53 | -1.53 |
| Tb927.1.4910:mRNA | expression site-associated gene 1 (ESAG1) protein | 1.782E-04 | 1.51 | -1.51 |
| Tb11.0290:mRNA | chrXI additional | 2.316E-02 | 1.31 | -1.31 |
| Tb927.7.410:mRNA | hypothetical protein | 2.655E-02 | 1.16 | -1.16 |
| Tb927.2.3310:mRNA | 65 kDa invariant surface glycoprotein | 3.022E-02 | 1.15 | -1.15 |
| Tb927.4.3550:mRNA | 60S ribosomal protein L13a | 4.148E-02 | 1.04 | -1.04 |
| Tb927.6.182:rRNA | M6 ribosomal RNA | 3.369E-21 | -5.36 | 5.36 |
| Tb927.4.200:mRNA | retrotransposon hot spot protein 1 (RHS1) | 4.492E-08 | -2.96 | 2.96 |
| Tb927.1.5170:mRNA | variant surface glycoprotein (VSG)-related | 8.914E-06 | -2.00 | 2.00 |
| Tb927.3.5820:pseudogenic_transcript | expression site-associated gene 11 (ESAG11) | 1.185E-05 | -3.55 | 3.55 |
| Tb11.v5.0301.1 | glycerol uptake protein | 7.464E-05 | -1.54 | 1.54 |
| Tb927.7.7540:pseudogenic_transcript | leucine-rich repeat protein 1 (LRRP1) | 7.685E-04 | -1.35 | 1.35 |
| Tb927.8.7830:mRNA | hypothetical protein | 9.703E-04 | -1.28 | 1.28 |
| Tb927.11.17550:mRNA | Unknown | 1.601E-03 | -4.55 | 4.55 |
| Tb927.10.2990:mRNA | nuclear cap binding complex subunit CBP110 | 1.601E-03 | -1.14 | 1.14 |
| Tb08.27P2.70:mRNA | Unknown | 1.798E-03 | -5.33 | 5.33 |
| Tb08.27P2.80:mRNA | hypothetical protein | 1.798E-03 | -4.24 | 4.24 |
| Tb927.1.5110:mRNA | expression site-associated gene 11 (ESAG11) protein | 2.501E-03 | -2.03 | 2.03 |
| Tb927.6.550:mRNA | hypothetical protein | 2.780E-03 | -1.58 | 1.58 |
| Tb11.v5.0376.1 | Trypanosomal VSG domain containing protein | 3.462E-03 | -2.72 | 2.72 |
| Tb927.11.17540:pseudogenic_transcript | variant surface glycoprotein (VSG) | 3.462E-03 | -2.69 | 2.69 |
| Tb11.v5.0381.1 | hypothetical protein | 3.462E-03 | -1.74 | 1.74 |
| Tb10.v4.0061:pseudogenic_transcript | Unknown | 6.156E-03 | -5.85 | 5.85 |
| Tb927.10.10220:mRNA | Unknown | 6.302E-03 | -3.24 | 3.24 |
| Tb927.9.15560:mRNA | BARP protein | 7.910E-03 | -2.65 | 2.65 |
| Tb11.v5.1039.1 | nuclear cap binding complex subunit CBP110 | 8.036E-03 | -1.08 | 1.08 |
| Tb927.10.12750:mRNA | hypothetical protein | 1.269E-02 | -1.59 | 1.59 |
| Tb927.4.240:pseudogenic_transcript | retrotransposon hot spot protein 4 (RHS4) | 1.563E-02 | -1.94 | 1.94 |
| Tb927.7.3830:mRNA | kinesin K39 | 1.728E-02 | -1.38 | 1.38 |
| Tb927.4.5350:pseudogenic_transcript | 3-methylcrotonyl-CoA carboxylase | 2.286E-02 | -1.08 | 1.08 |
| Tb927.11.810:mRNA | hypothetical protein | 2.286E-02 | -1.05 | 1.05 |
| Tb927.1.5240:mRNA | Unknown | 2.308E-02 | -4.80 | 4.80 |
| Tb10.v4.0070:mRNA | Unknown | 2.405E-02 | -4.67 | 4.67 |
| Tb927.2.240:mRNA | retrotransposon hot spot protein 5 (RHS5) | 2.613E-02 | -1.30 | 1.30 |
| Tb10.v4.0096:mRNA | variant surface glycoprotein (VSG) | 3.139E-02 | -2.43 | 2.43 |
| Tb927.10.11470:mRNA | hypothetical protein | 3.264E-02 | -1.01 | 1.01 |
| Tb09.v4.0143:pseudogenic_transcript | variant surface glycoprotein (VSG) | 3.276E-02 | -1.85 | 1.85 |
| Tb09.v4.0109:mRNA | hypothetical protein | 3.276E-02 | -1.66 | 1.66 |
| Tb927.1.2880:mRNA | pteridine transporter | 3.286E-02 | -1.30 | 1.30 |
| Tb927.11.17460:mRNA | variant surface glycoprotein (VSG) | 4.090E-02 | -2.47 | 2.47 |
| Tb927.1.4870:mRNA | expression site-associated gene 1 (ESAG1) protein | 4.148E-02 | -1.21 | 1.21 |
| Tb11.1060:pseudogenic_transcript | Unknown | 4.162E-02 | -4.52 | 4.52 |
| Tb927.7.6540:mRNA | variant surface glycoprotein (VSG) | 4.162E-02 | -2.29 | 2.29 |
| Tb11.1470:pseudogenic_transcript | variant surface glycoprotein (VSG) | 4.162E-02 | -1.80 | 1.80 |
| Tb927.6.5320:pseudogenic_transcript | Unknown | 4.264E-02 | -4.33 | 4.33 |
| Tb927.5.2360:mRNA | hypothetical protein | 4.444E-02 | -1.05 | 1.05 |
| Tb11.1200:mRNA | hypothetical protein | 4.607E-02 | -1.10 | 1.10 |
| Tb11.v5.0416.1 | Variant Surface Glycoprotein | 4.947E-02 | -1.79 | 1.79 |
| Tb927.7.6780:mRNA | hypothetical protein | 4.947E-02 | -1.13 | 1.13 |
| Tb927.11.10740:mRNA | vacuolar sorting-associated protein-like | 4.607E-02 | 0.95 | 0.95 |
| Tb927.10.560:mRNA | 40S ribosomal proteins S11 | 4.148E-02 | 0.91 | 0.91 |
| Tb927.5.620:mRNA | invariant surface glycoprotein | 2.892E-02 | 0.91 | 0.91 |
| Tb11.v5.0871.1 | hypothetical protein | 2.069E-02 | 0.89 | 0.89 |
| Tb927.9.14000:mRNA | 60S ribosomal protein L12 | 2.344E-02 | 0.80 | 0.80 |
| Tb11.v5.0326.1 | retrotransposon hot spot (RHS) protein | 4.148E-02 | 0.74 | 0.74 |
| Tb927.3.1290:mRNA | cullin 4B | 4.148E-02 | -0.68 | -0.68 |
| Tb927.1.180:mRNA | retrotransposon hot spot protein 1 (RHS1) | 4.325E-02 | -0.70 | -0.70 |
| Tb927.11.16010:mRNA | hypothetical protein | 4.325E-02 | -0.74 | -0.74 |
| Tb927.11.4280:mRNA | hypothetical protein | 4.264E-02 | -0.76 | -0.76 |
| Tb927.9.12160:mRNA | hypothetical protein | 2.286E-02 | -0.78 | -0.78 |
| Tb927.11.890:mRNA | hypothetical protein | 1.394E-02 | -0.78 | -0.78 |
| Tb927.8.6790:mRNA | hypothetical protein | 1.149E-02 | -0.79 | -0.79 |
| Tb927.1.220:mRNA | retrotransposon hot spot protein 1 (RHS1) | 1.394E-02 | -0.81 | -0.81 |
| Tb927.1.120:mRNA | retrotransposon hot spot protein 4 (RHS4) | 3.324E-02 | -0.84 | -0.84 |
| Tb927.10.14510:mRNA | root hair defective 3 GTP-binding protein (RHD3) | 4.461E-03 | -0.84 | -0.84 |
| Tb927.10.14900:mRNA | hypothetical protein | 4.739E-02 | -0.85 | -0.85 |
| Tb927.9.5520:mRNA | ubiquitin carboxyl-terminal hydrolase | 6.156E-03 | -0.86 | -0.86 |
| Tb927.7.6650:mRNA | Colon cancer-associated protein Mic1-like | 3.286E-02 | -0.86 | -0.86 |
| Tb927.11.6120:mRNA | ABC transporter | 2.640E-02 | -0.87 | -0.87 |
| Tb927.7.6760.1:mRNA | hypothetical protein | 9.459E-03 | -0.87 | -0.87 |
| Tb927.2.5480:mRNA | hypothetical protein | 4.577E-02 | -0.92 | -0.92 |
| Tb927.10.2570:mRNA | lysosomal alpha-mannosidase precursor | 2.292E-03 | -0.93 | -0.93 |
| Tb11.02.5130b.1 | neurobeachin/beige protein | 4.148E-02 | -0.94 | -0.94 |
| Tb927.4.4510:mRNA | protein phosphatase 2C | 4.148E-02 | -0.98 | -0.98 |
| Tb927.1.5150:mRNA | hypothetical protein | 2.501E-03 | -0.99 | -0.99 |

**Table S2:** Significant (p<0.05) gene ontology enrichment of *T.b. rhodesiense* biological processes of differentially enriched genes (DEGs) that were upregulated (log2FC > 1) in Nkhotakota focus and loaded in TritrypDB. The fold enrichment is the percentage of genes loaded divide by the percentage of genes with this term in the background. The p-value measured the Fishers exact test.

| Gene Ontology | Biological Process | Fold Enrichment (FE) | P-value of FE |
| --- | --- | --- | --- |
| GO:0010256 | Endomembrane system organization | 52.9 | 2.09E-05 |
| GO:0006890 | Retrograde vesicle-mediated transport, Golgi to endoplasmic reticulum | 80.32 | 0.000265 |
| GO:0007030 | Golgi organization | 49.86 | 0.000698 |
| GO:0048193 | Golgi vesicle transport | 16.06 | 0.006566 |
| GO:0007009 | Plasma membrane organization | 103.27 | 0.009646 |
| GO:0046907 | Intracellular transport | 5.71 | 0.013654 |
| GO:0033036 | macromolecule localization | 5.63 | 0.014146 |
| GO:0051649 | establishment of localization in cell | 5.6 | 0.014345 |
| GO:0071850 | mitotic cell cycle arrest | 60.24 | 0.016484 |
| GO:0071705 | nitrogen compound transport | 5.21 | 0.017424 |
| GO:0009299 | mRNA transcription | 51.64 | 0.019207 |
| GO:0042789 | mRNA transcription by RNA polymerase II | 51.64 | 0.019207 |
| GO:0051641 | cellular localization | 4.99 | 0.019631 |
| GO:0016043 | cellular component organization | 3.52 | 0.021239 |
| GO:0071702 | organic substance transport | 4.82 | 0.021481 |
| GO:0000028 | ribosomal small subunit assembly | 40.16 | 0.024632 |
| GO:0034249 | negative regulation of cellular amide metabolic process | 34.42 | 0.028683 |
| GO:0017148 | negative regulation of translation | 34.42 | 0.028683 |
| GO:0006406 | mRNA export from nucleus | 26.77 | 0.036739 |
| GO:0051248 | negative regulation of protein metabolic process | 25.82 | 0.038075 |
| GO:0032269 | negative regulation of cellular protein metabolic process | 25.82 | 0.038075 |
| GO:0016192 | vesicle-mediated transport | 6.29 | 0.038778 |
| GO:0071840 | cellular component organization or biogenesis | 2.92 | 0.039403 |
| GO:0006886 | intracellular protein transport | 6.15 | 0.040331 |
| GO:0000462 | maturation of SSU-rRNA from tricistronic rRNA transcript (SSU-rRNA, 5.8S rRNA, LSU-rRNA) | 24.1 | 0.040744 |
| GO:0051028 | mRNA transport | 23.32 | 0.042075 |
| GO:0034645 | cellular macromolecule biosynthetic process | 3.68 | 0.04321 |
| GO:0009059 | macromolecule biosynthetic process | 3.66 | 0.04377 |
| GO:0006405 | RNA export from nucleus | 21.26 | 0.04606 |

**Tables S3:**
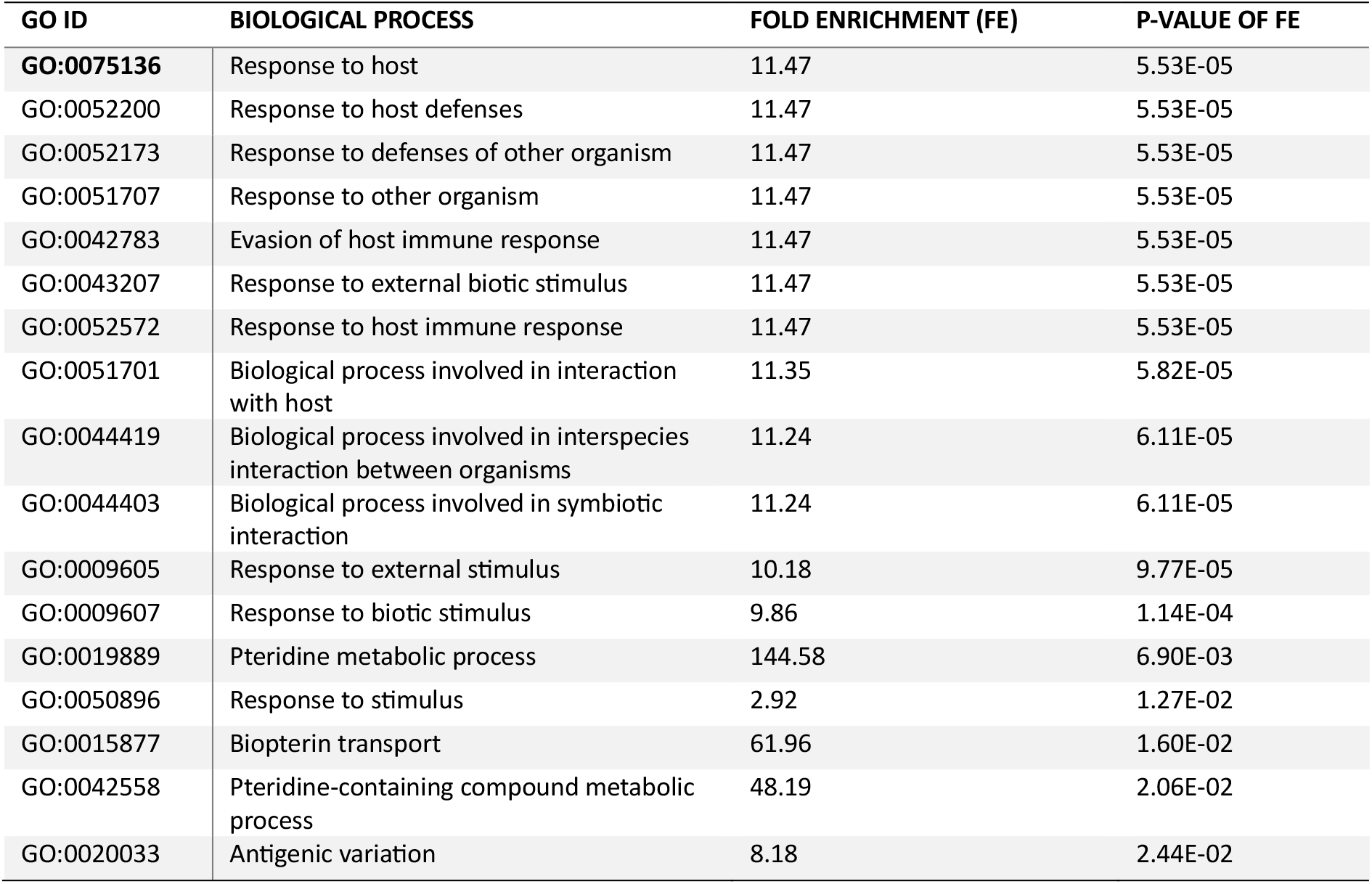
Significant (p<0.05) gene ontology (GO) enrichment of T.b. rhodesiense biological processes of differentially enriched genes (DEGs) that were upregulated (log2FC > 1) in Rumphi focus and loaded in TritrypDB. The fold enrichment is the percentage of genes loaded divide by the percentage of genes with this term in the background. The p-value measured the Fishers exact test.

## References

1. WHO. Report of the third WHO stakeholders meeting on rhodesiense human African trypanosomiasis, Geneva, Switzerland, 10–11 April 2019.: WHO; 2020 [74]. Available from: https://www.who.int/publications/i/item/9789240012936.

2. Nambala P, Mulindwa J, Chammudzi P, Senga E, Lemelani M, Zgambo D, et al. Persistently High Incidences of Trypanosoma brucei rhodesiense Sleeping Sickness With Contrasting Focus-Dependent Clinical Phenotypes in Malawi. Front Trop Dis. 2022:3:824484.

3. Gibson W, Peacock L, Ferris V, Fischer K, Livingstone J, Thomas J, et al. Genetic recombination between human and animal parasites creates novel strains of human pathogen. PLoS Negl Trop Dis. 2015;9(3):e0003665.

4. Duffy CW, MacLean L, Sweeney L, Cooper A, Turner CM, Tait A, et al. Population genetics of Trypanosoma brucei rhodesiense: clonality and diversity within and between foci. PLoS Negl Trop Dis. 2013;7(11):e2526.

5. MacLean L, Chisi JE, Odiit M, Gibson WC, Ferris V, Picozzi K, et al. Severity of human african trypanosomiasis in East Africa is associated with geographic location, parasite genotype, and host inflammatory cytokine response profile. Infect Immun. 2004;72(12):7040–4.

6. Echodu R, Sistrom M, Bateta R, Murilla G, Okedi L, Aksoy S, et al. Genetic diversity and population structure of Trypanosoma brucei in Uganda: implications for the epidemiology of sleeping sickness and Nagana. PLoS Negl Trop Dis. 2015;9(2):e0003353.

7. Nambala P, Mulindwa J, Noyes H, Namulondo J, Nyangiri O, Matovu E, et al. Distinct Differences in Gene Expression Profiles in Early and Late Stage Rhodesiense HAT Individuals in Malawi. bioRxiv; 2022.

8. Mathieu-Daude F, Welsh J, Davis C, McClelland M. Differentially expressed genes in the Trypanosoma brucei life cycle identified by RNA fingerprinting. Mol Biochem Parasitol. 1998;92(1):15–28.

9. Monk SL, Simmonds P, Matthews KR. A short bifunctional element operates to positively or negatively regulate ESAG9 expression in different developmental forms of Trypanosoma brucei. J Cell Sci. 2013;126(Pt 10):2294–304.

10. Barnwell EM, van Deursen FJ, Jeacock L, Smith KA, Maizels RM, Acosta-Serrano A, et al. Developmental regulation and extracellular release of a VSG expression-site-associated gene product from Trypanosoma brucei bloodstream forms. J Cell Sci. 2010;123(Pt 19):3401–11.

11. Guther ML, Beattie K, Lamont DJ, James J, Prescott AR, Ferguson MA. Fate of glycosylphosphatidylinositol (GPI)-less procyclin and characterization of sialylated non-GPI-anchored surface coat molecules of procyclic-form Trypanosoma brucei. Eukaryot Cell. 2009;8(9):1407–17.

12. Guther ML, Lee S, Tetley L, Acosta-Serrano A, Ferguson MA. GPI-anchored proteins and free GPI glycolipids of procyclic form Trypanosoma brucei are nonessential for growth, are required for colonization of the tsetse fly, and are not the only components of the surface coat. Mol Biol Cell. 2006;17(12):5265–74.

13. Aslett M, Aurrecoechea C, Berriman M, Brestelli J, Brunk BP, Carrington M, et al. TriTrypDB: a functional genomic resource for the Trypanosomatidae. Nucleic Acids Res. 2010;38(Database issue):D457–62.

14. Ali I, Yang WC. The functions of kinesin and kinesin-related proteins in eukaryotes. Cell Adh Migr. 2020;14(1):139–52.

15. Gerald NJ, Coppens I, Dwyer DM. Molecular dissection and expression of the LdK39 kinesin in the human pathogen, Leishmania donovani. Mol Microbiol. 2007;63(4):962–79.

16. Supek F, Bosnjak M, Skunca N, Smuc T. REVIGO summarizes and visualizes long lists of gene ontology terms. PLoS One. 2011;6(7):e21800.

17. Sudipta Hazra SG, Banasri Hazra. Phytochemicals With Antileishmanial Activity: Prospective Drug Targets. Studies in Natural Products Chemistry. 522017. p. 303–36.

18. MacLean LM, Odiit M, Chisi JE, Kennedy PG, Sternberg JM. Focus-specific clinical profiles in human African Trypanosomiasis caused by Trypanosoma brucei rhodesiense. PLoS Negl Trop Dis. 2010;4(12):e906.

19. Mulindwa J, Leiss K, Ibberson D, Kamanyi Marucha K, Helbig C, Melo do Nascimento L, et al. Transcriptomes of Trypanosoma brucei rhodesiense from sleeping sickness patients, rodents and culture: Effects of strain, growth conditions and RNA preparation methods. PLoS Negl Trop Dis. 2018;12(2):e0006280.

20. Mulindwa J, Merce C, Matovu E, Enyaru J, Clayton C. Transcriptomes of newly-isolated Trypanosoma brucei rhodesiense reveal hundreds of mRNAs that are co-regulated with stumpy-form markers. BMC Genomics. 2015;16:1118.

21. Holsinger KE, Weir BS. Genetics in geographically structured populations: defining, estimating and interpreting F(ST). Nat Rev Genet. 2009;10(9):639–50.

22. Willing EM, Dreyer C, van Oosterhout C. Estimates of genetic differentiation measured by F(ST) do not necessarily require large sample sizes when using many SNP markers. PLoS One. 2012;7(8):e42649.

23. Burns JM, Jr., Shreffler WG, Benson DR, Ghalib HW, Badaro R, Reed SG. Molecular characterization of a kinesin-related antigen of Leishmania chagasi that detects specific antibody in African and American visceral leishmaniasis. Proc Natl Acad Sci U S A. 1993;90(2):775–9.

24. Chisi J, Nkhoma A, Sternberg J. Presentation of trypanosomiasis in nkhotakota. Malawi Med J. 2007;19(4):140–1.

25. Rojas F, Silvester E, Young J, Milne R, Tettey M, Houston DR, et al. Oligopeptide Signaling through TbGPR89 Drives Trypanosome Quorum Sensing. Cell. 2018.

26. Jayaraman S, Harris C, Paxton E, Donachie AM, Vaikkinen H, McCulloch R, et al. Application of long read sequencing to determine expressed antigen diversity in Trypanosoma brucei infections. PLoS Negl Trop Dis. 2019;13(4):e0007262.

27. Autheman D, Crosnier C, Clare S, Goulding DA, Brandt C, Harcourt K, et al. An invariant Trypanosoma vivax vaccine antigen induces protective immunity. Nature. 2021;595(7865):96–100.

28. Radwanska M, Chamekh M, Vanhamme L, Claes F, Magez S, Magnus E, et al. The serum resistance-associated gene as a diagnostic tool for the detection of Trypanosoma brucei rhodesiense. Am J Trop Med Hyg. 2002;67(6):684–90.

29. Mulindwa J, Fadda A, Merce C, Matovu E, Enyaru J, Clayton C. Methods to determine the transcriptomes of trypanosomes in mixtures with mammalian cells: the effects of parasite purification and selective cDNA amplification. PLoS Negl Trop Dis. 2014;8(4):e2806.

30. Love MI, Huber W, Anders S. Moderated estimation of fold change and dispersion for RNA-seq data with DESeq2. Genome Biol. 2014;15(12):550.

31. So J, Sudlow S, Sayeed A, Grudda T, Deborggraeve S, Ngoyi DM, et al. VSGs Expressed during Natural T. b. gambiense Infection Exhibit Extensive Sequence Divergence and a Subspecies-Specific Bias towards Type B N-Terminal Domains. mBio. 2022;13(6):e0255322.

32. Holsinger KE. Next generation population genetics and phylogeography. Mol Ecol. 2010;19(12):2361–3.

33. Dessau RB, Pipper CB. [’’R"--project for statistical computing]. Ugeskr Laeger. 2008;170(5):328–30.

